# Tissue-resolved photosynthetic responses in living leaves revealed by microscopic imaging-pulse-amplitude-modulation

**DOI:** 10.64898/2026.09.09.750010

**Authors:** Yuta Kato, Ko Takeuchi, Shintaro Harimoto, Koich Kobayashi, Kentaro Ifuku

## Abstract

Leaves of terrestrial plants possess heterogeneous anatomical structures composed of multiple cell layers. Although biochemical differences among leaf tissues have been inferred from protein analyses and anatomical studies, direct comparisons of dynamic photochemical responses among tissues while preserving their spatial context remain challenging. Leaves experience intrinsically heterogeneous environments because incident light enters primarily from above and propagates through complex internal leaf structures. Therefore, analyzing photosynthetic activity at the tissue and cellular levels is essential for understanding how photosynthesis operates within structurally heterogeneous leaves.

Here, we combined live leaf-section imaging with microscopic Imaging-PAM chlorophyll fluorescence measurements to analyze photochemical responses at the tissue level in living leaf sections. In dorsiventral dicot leaves, palisade tissues exhibited a higher effective PSII quantum yield [Y(II)] and more rapid induction of regulated energy dissipation [Y(NPQ)] than spongy tissues, indicating higher photosynthetic capacity and photoprotective activity. In contrast, rice leaves, which lack palisade–spongy differentiation, showed uniform photochemical responses along the adaxial–abaxial axis. In the C_4_ plant finger millet, mesophyll and bundle sheath cells displayed distinct photochemical responses consistent with their functional differentiation in C_4_ photosynthesis.

These results demonstrate that photosynthetic responses are spatially organized according to leaf anatomical structure. Although section-based measurements do not reproduce the native optical and gas environments of intact leaves, they enable the comparison of intrinsic tissue-specific photochemical properties under approximately equivalent illumination. The present approach enables tissue-resolved chlorophyll fluorescence analysis in living leaves, providing a new framework for investigating how leaf architecture shapes the spatial organization of photosynthetic activity.

## Introduction

Leaves have heterogeneous internal structures. In many plant species, leaves are anatomically differentiated into palisade tissues on the adaxial side and spongy tissues on the abaxial side. This organization reportedly facilitates the efficient utilization of incident light entering from the adaxial surface (Vogelmann and Martin 1993; Vogelmann and Evans 2002; Terashima et al. 2009). When light is irradiated from the adaxial side, its intensity progressively attenuates toward the abaxial side, resulting in a steep internal light gradient along the adaxial–abaxial axis (Takahashi et al. 1994; Vogelmann and Evans 2002; Terashima et al. 2016). Consistent with this gradient, several studies have reported that chlorophyll content, Rubisco abundance, and photosynthetic electron transport components are higher in tissues located closer to the adaxial surface than in those closer to the abaxial side (Terashima and Inoue 1984; 1985a, b). Additionally, leaf anatomical organization is not limited to vertical differentiation such as palisade and spongy tissues. Some monocots, including rice, possess mesophyll tissues composed of lobed cells without distinct palisade and spongy layers (Chonan 1967; Oi et al. 2017). In C_4_ plants, leaf structure is radially organized around the vascular bundles, with functionally distinct mesophyll and bundle sheath cells (Brown 1975; Dengler and Nelson 1999). Leaf anatomical organization thus reflects not only adaptation to vertical light gradients but also variation in photosynthetic function among plant species, leading to diverse structural arrangements.

Despite this recognition of internal light gradients and anatomical heterogeneity, how individual tissues in a living leaf dynamically respond to light remains poorly understood. This difficulty arises largely from the strong scattering properties of leaf tissues, which cause light signals to be spatially integrated across the leaf thickness in conventional measurements, making it difficult to distinguish signals originating from different tissue layers. Conversely, biochemical analyses of physically separated tissues disrupt their spatial context and are unable to capture real-time photochemical regulation under physiologically active conditions. Consequently, most current knowledge of tissue-specific photosynthetic function is based on static biochemical composition, anatomical observations, or modeling, rather than direct measurements of photosynthetic activity at the tissue level in intact living leaves (Terashima et al. 2016, Xiao et al. 2016).

Chlorophyll fluorescence analysis using pulse-amplitude-modulation (PAM) fluorometry is a standard method for assessing photosynthetic activity (Klughammer and Schreiber 2008; Baker 2008). Conventional PAM fluorometry, however, illuminates a leaf from one side and detects spatially integrated fluorescence. Measurements from the adaxial or abaxial surfaces can partially bias the contribution of palisade and spongy tissues, respectively, but the relative contribution of each tissue remains uncertain (Peguero-Pina et al. 2009; Oguchi et al. 2011). An alternative and innovative approach using fiber-optic PAM has been applied to obtain chlorophyll fluorescence signals at specific depths within leaves by inserting a microprobe into the tissue (Schreiber et al. 1996; Oguchi et al. 2011). These pioneering studies have provided important insights into vertical heterogeneity of photochemical responses within leaves. However, because measurements are obtained at one location at a time, spatial coverage remains limited and simultaneous multi-point observations are difficult. In addition, practical constraints related to leaf thickness and rigidity restrict the applicability of this approach across diverse plant species. Chlorophyll fluorescence measurements of leaf cross-sections have also been reported, particularly in C_4_ species (Liu et al. 2022; Ouyang et al. 2024), but their broader use for comparative tissue physiology remains limited. Previously, we established a live leaf-section imaging method enabling long-term observation of chloroplast dynamics in excised leaves (Kato et al. 2025a, b). By combining this method with microscopic Imaging-PAM, the present study enables multiple tissues to be simultaneously analyzed at defined spatial positions under a common illumination regime.

In this study, we investigated whether different forms of leaf anatomical specialization are accompanied by distinct spatial patterns of photochemical regulation. We first compared palisade and spongy tissues in dorsiventral dicot leaves, which develop under contrasting internal light environments. We then examined rice mesophyll tissue, which lacks distinct palisade–spongy differentiation, to test whether its photochemical responses are more uniform along the adaxial–abaxial axis. Finally, we examined whether C_4_ leaves exhibit functional partitioning of photosynthetic reactions between mesophyll (M) and bundle sheath (BS) cells. Through these comparisons, we aimed to reveal the anatomy-dependent photochemical architecture of leaves across different plant species.

## Results and Discussion

The present study demonstrates that different tissues within a leaf exhibit distinct photochemical responses to the same light stimulus from the sectioning face. The heterogeneous internal light environment and anatomical differentiation of leaves have long been recognized (Terashima and Saeki, 1983; Vogelmann and Evans, 2002; Terashima et al., 2016). Our results provide direct functional evidence at the tissue level in living leaf sections under equivalent illumination conditions, supporting these long-standing concepts. In the present study, the induction phase (dark-to-light transitions) of chlorophyll fluorescence was monitored under actinic light (AL) (115.8 μmol photons m^-2^ s^-1^) for 9.5 min after 15 min of dark adaptation. It should be noted that, during the slow induction phase of chlorophyll fluorescence, CO_2_ availability, which may vary among different tissues in intact leaves because of their heterogeneous internal structures, can influence both photochemical and non-photochemical yields. However, in the present study, the measurements were performed after infiltration with 2 mM NaHCO_3_, a treatment that would be expected to reduce spatial heterogeneity in CO_2_ availability across the leaf section.

### Palisade and spongy tissues in dicots

In cucumber leaves, Y(II) (the effective quantum yield of PSII under light illumination) increased rapidly until approximately 90–120 s, reaching approximately 0.25 in the palisade tissue and approximately 0.15 in the spongy tissue. Thereafter, Y(II) continued to increase gradually in both tissues (Fig. 2, Supplementary Movie 1). On the other hand, Y(NPQ) (regulated energy dissipation in PSII) increased more rapidly in the palisade tissue than in the spongy tissue immediately after AL illumination, but Y(NPQ) in the spongy tissue gradually approached that of the palisade tissue, with both tissues eventually reaching approximately 0.3. Y(NO) (nonregulated energy dissipation in PSII) decreased rapidly after AL irradiation, then declined more gradually, reaching approximately 0.4 in the palisade tissue and approximately 0.5 in the spongy tissue. There was a significant difference in F_v_/F_m_ between the palisade and the spongy tissues (Table 1). In addition, measurements using fiber-type PAM (Dual-PAM 100) on leaf discs revealed that Y(II) was higher when measured from the adaxial side than from the abaxial side (Supplementary Fig. S2). The maximum value of Y(NPQ) was also higher when measurements were performed from the adaxial surface.

**Fig. 1.**
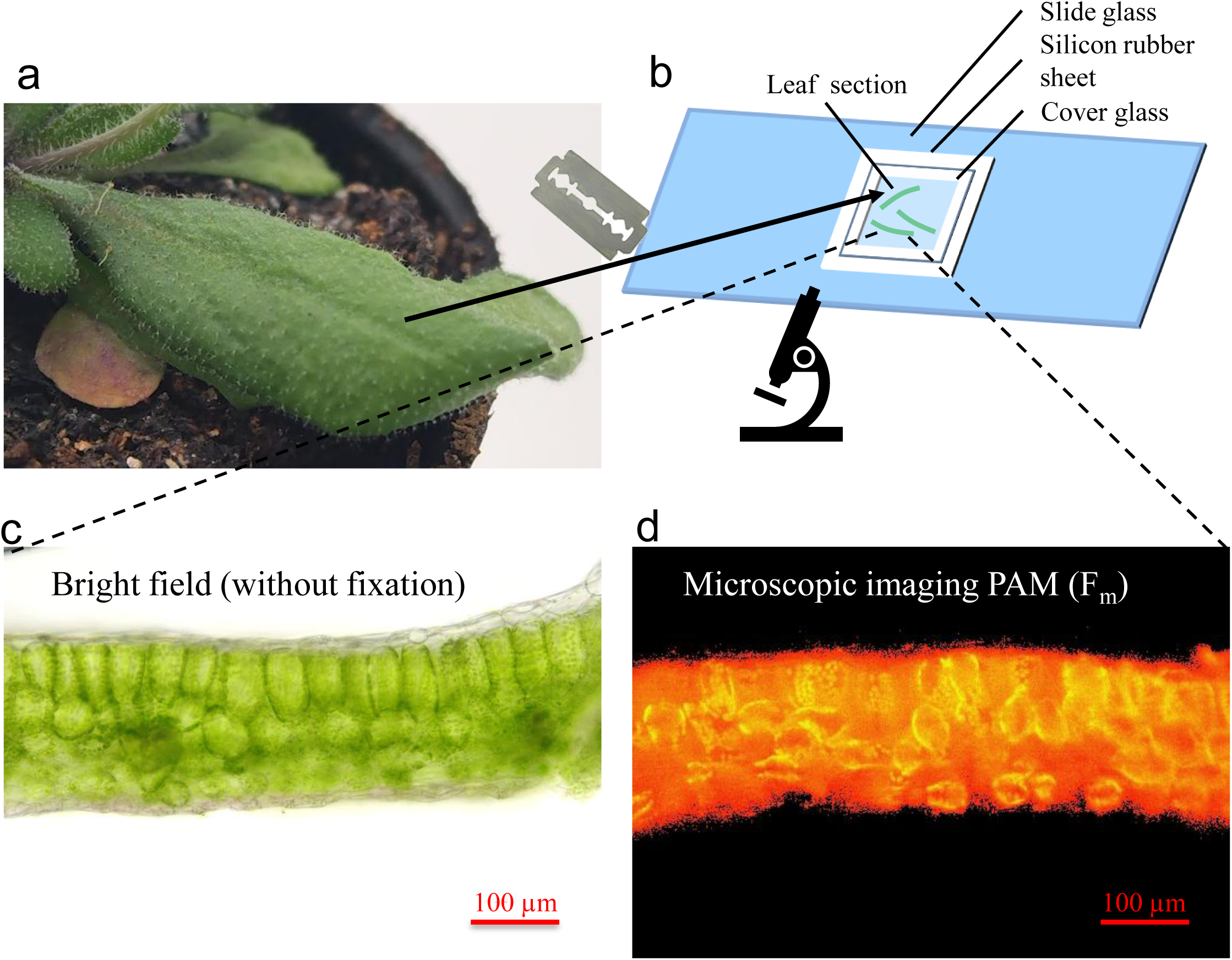
Preparation and microscopic Imaging-PAM analysis of live leaf sections. (a) *Arabidopsis thaliana* leaf. (b) Schematic diagram of the live leaf-section mount. (c) Bright-field microscopic image of a transverse leaf section of *Arabidopsis thaliana*. (d) Fm image of a transverse leaf section of *Arabidopsis thaliana* acquired using microscopic Imaging-PAM. The upper side of the microscopic image corresponds to the adaxial side of the leaf.

**Fig. 2.**
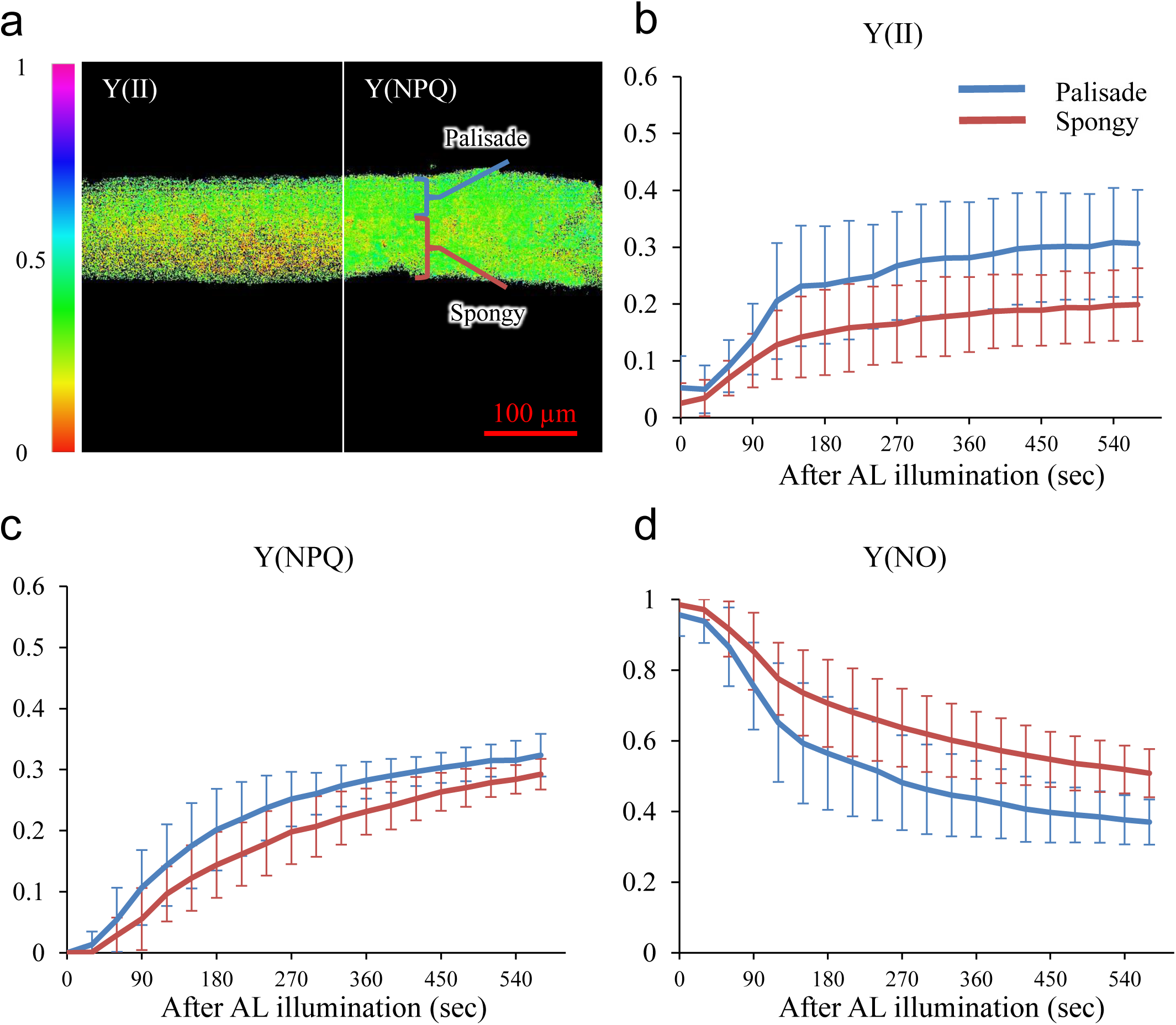
Microscopic Imaging-PAM analysis of cucumber leaf transverse sections. Chlorophyll fluorescence during the induction phase of photosynthesis was measured in dark-adapted leaves using microscopic Imaging-PAM at room temperature under actinic light (AL; PPFD, 115.8 µmol m⁻² s⁻¹) for 9.5 min. (a) Y(II) (left side) and Y(NPQ) (right side) images of a transverse leaf section of cucumber acquired using microscopic Imaging-PAM at the end of the induction period. The upper side corresponds to the adaxial side of the leaf. (b–d) Induction curves of Y(II), Y(NPQ), and Y(NO) measured in palisade and spongy tissues. Values are means ± SD. n = 3.

**Table 1.**
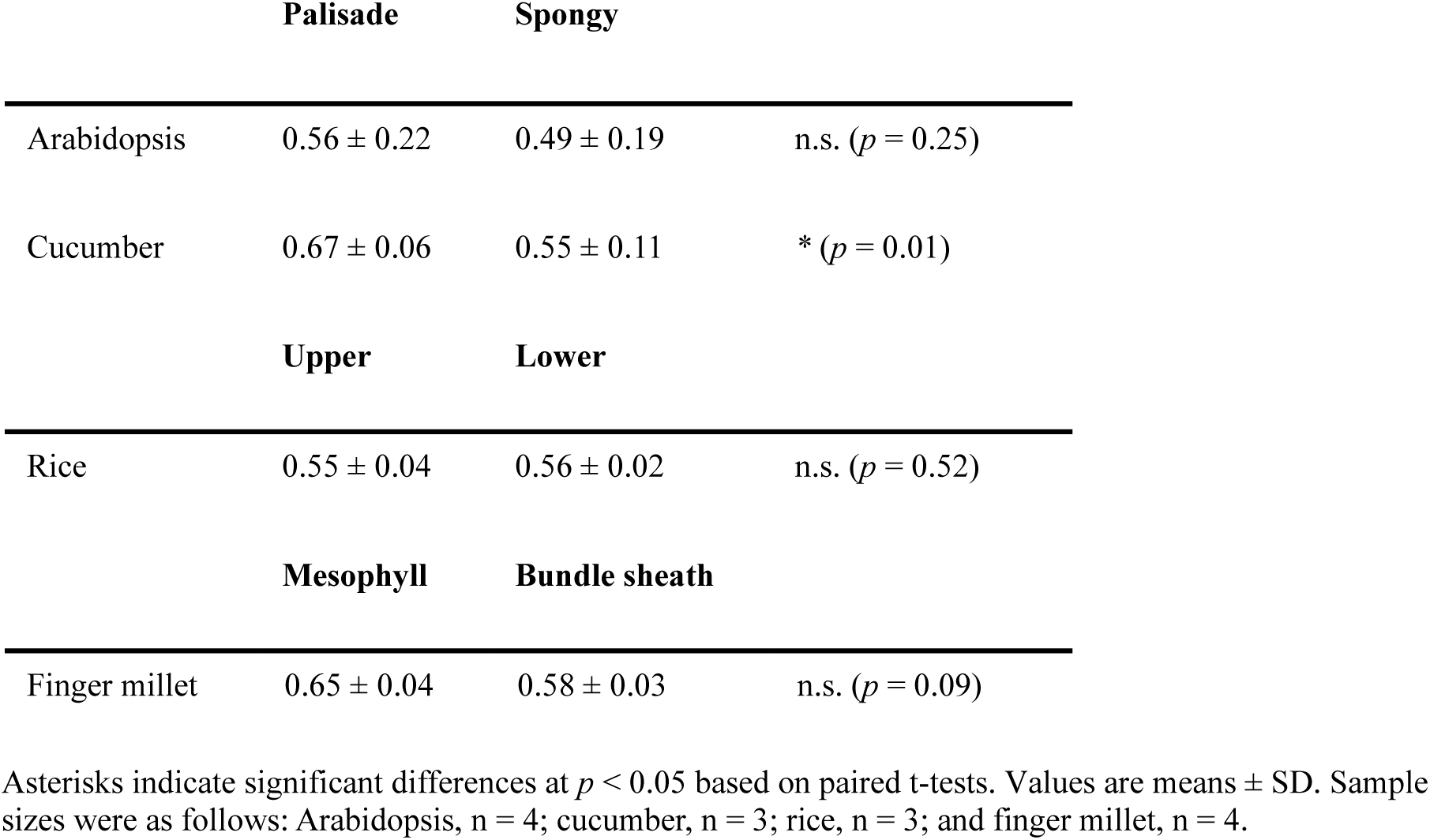
Fv/Fm values in each region.

|  | **Palisade** | **Spongy** |  |
| --- | --- | --- | --- |
| Arabidopsis | 0.56 ± 0.22 | 0.49 ± 0.19 | n.s. ($p = 0.25$) |
| Cucumber | 0.67 ± 0.06 | 0.55 ± 0.11 | * ($p = 0.01$) |
|  | **Upper** | **Lower** |  |
| Rice | 0.55 ± 0.04 | 0.56 ± 0.02 | n.s. ($p = 0.52$) |
|  | **Mesophyll** | **Bundle sheath** |  |
| Finger millet | 0.65 ± 0.04 | 0.58 ± 0.03 | n.s. ($p = 0.09$) |

Asterisks indicate significant differences at $p < 0.05$ based on paired t-tests. Values are means ± SD. Sample sizes were as follows: Arabidopsis, n = 4; cucumber, n = 3; rice, n = 3; and finger millet, n = 4.

In *Arabidopsis* leaves, Y(II) increased rapidly in both palisade and spongy mesophyll tissues and reached an approximately steady state within approximately 120–150 s (Fig. 3, Supplementary Movie 2). The steady-state Y(II) was higher in the palisade tissue (approximately 0.4) than in the spongy tissue (approximately 0.3). Y(NPQ) showed a rapid increase in both tissues within 30–60 s, followed by a transient decline and a subsequent gradual increase. The initial increase in Y(NPQ) was higher in the palisade tissue (approximately 0.3) than in the spongy tissue (approximately 0.2). Y(NO) decreased rapidly within 60 s and remained slightly lower thereafter in the palisade tissue than in the spongy tissues. There was no significant difference in F_v_/F_m_ between the palisade and the spongy tissues (Table 1).

**Fig. 3.**
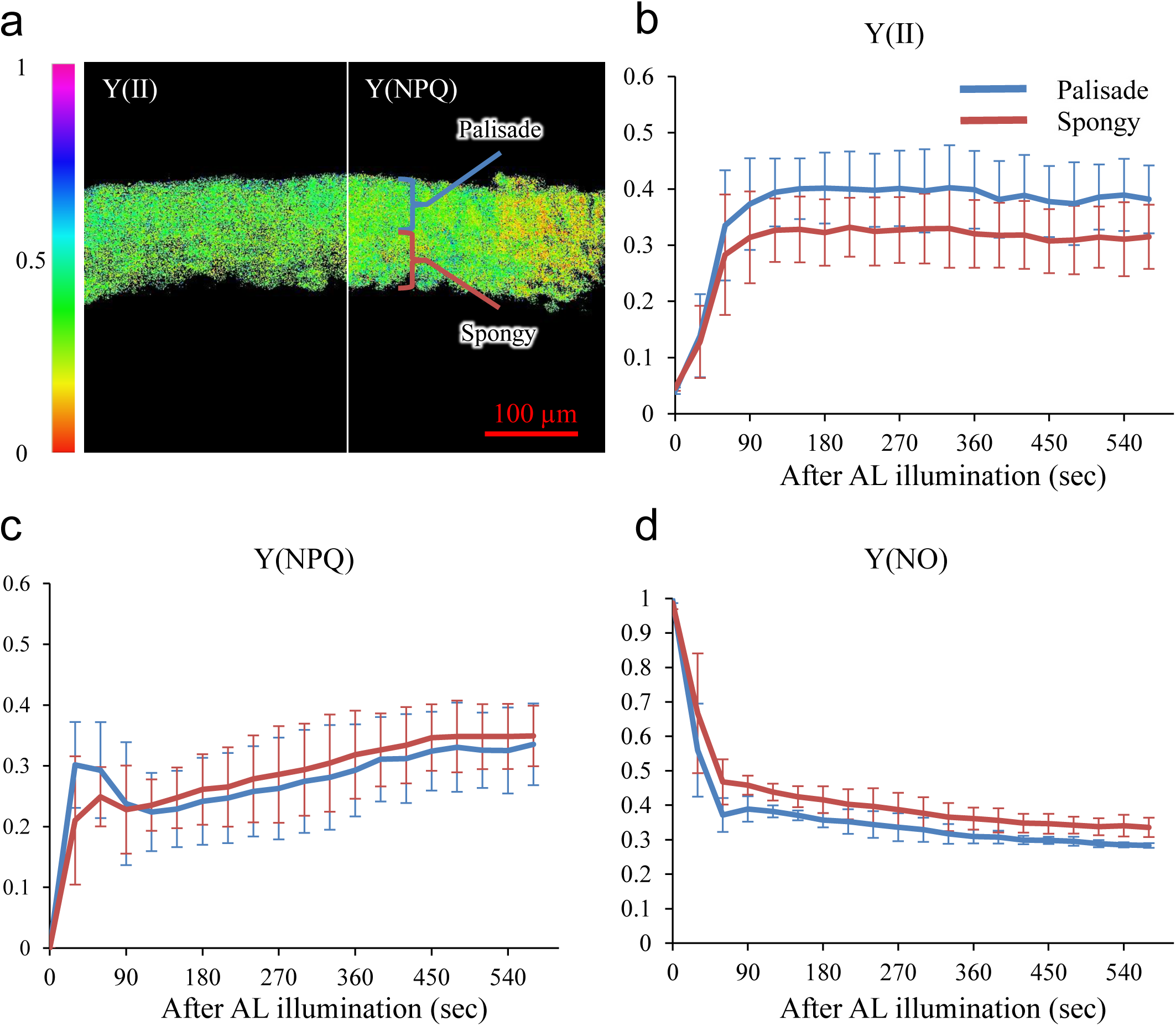
Microscopic Imaging-PAM analysis of Arabidopsis leaf transverse sections. (a) Y(II) (left side) and Y(NPQ) (right side) images of a transverse leaf section of Arabidopsis at the end of the induction period. The upper side corresponds to the adaxial side of the leaf. (b–d) Y(II), Y(NPQ), and Y(NO) measured in palisade and spongy tissues. Values are means ± SD. n = 4.

Camellia leaves are thick and possess multiple layers of palisade and spongy mesophyll cells. Therefore, we divided the palisade tissue into three regions corresponding to the first, second, and third cell layers and the spongy tissue into three regions for measurement. Y(II) increased more rapidly and reached higher values in the palisade tissue layers located closer to the adaxial epidermis. Among the spongy tissues, Y(II) values were similar (Fig. 4, Supplementary Movie 3). Y(NPQ) increased more rapidly and reached higher values in the palisade and spongy tissues closer to the adaxial epidermis. Y(NO) decreased more rapidly and reached lower values in the palisade and spongy tissues closer to the adaxial epidermis. F_v_/F_m_ in the first palisade layer was significantly higher than that in the spongy tissues (Fig. 4E).

**Fig. 4.**
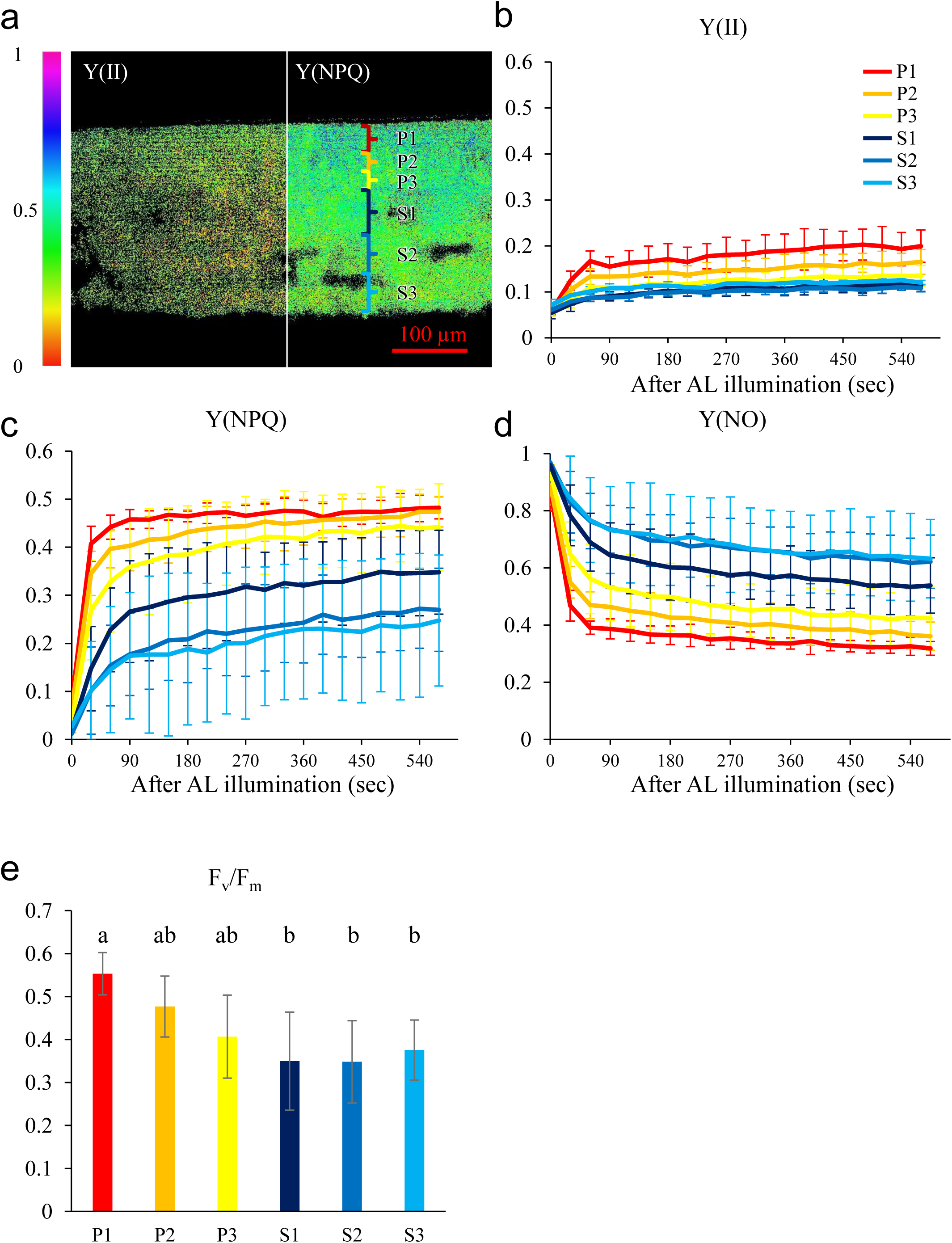
Microscopic Imaging-PAM analysis of camellia leaf transverse sections. (a) Y(II) (left side) and Y(NPQ) (right side) images of a transverse leaf section of camellia acquired using microscopic Imaging-PAM at the end of the induction period. The upper side corresponds to the adaxial side of the leaf. (b–d) Induction curves of Y(II), Y(NPQ), and Y(NO) measured in each region. Values are means ± SD. n = 5. (e) Fv/Fm values for each region. Different letters indicate significant differences (*p* < 0.05) according to the Tukey–Kramer test.

In dicot leaves possessing palisade and spongy tissues (*Arabidopsis*, cucumber, and camellia), palisade tissues consistently exhibited higher Y(II), while Y(NPQ) generally increased more rapidly in palisade tissues than in spongy tissues (Figs. 2–4). As the tissues were illuminated from the section surface, the palisade and spongy tissues were exposed to similar incident light during the measurement. Therefore, the observed differences are unlikely to result from differences in light quantity during the measurement, but rather reflect intrinsic tissue-specific differences. Previous studies have shown that palisade tissues contain more photosynthetic apparatus, including Rubisco and components of the photosynthetic electron transport chain, than spongy tissues (Terashima and Inoue, 1984; 1985a, b; Nishio et al., 1993). This higher photosynthetic capacity may contribute to the higher Y(II) in palisade tissues. The more rapid increase in Y(NPQ) in palisade tissues may reflect a higher capacity to induce energy-dependent quenching (qE), the fastest component of NPQ. qE depends on the formation of a proton gradient across the thylakoid membrane, PsbS, and xanthophyll-cycle activity (Zuo 2025), and its capacity is enhanced by acclimation to high light through increased PsbS abundance and expansion of the violaxanthin–antheraxanthin–zeaxanthin pool (Thayer and Björkman 1990; Demmig-Adams and Adams 2006; Zia et al. 2011). Higher carotenoid abundance has also been reported in palisade mesophyll exposed to stronger incident light (Nishio et al. 1993). Thus, the higher Y(II) and more rapid induction of Y(NPQ) in palisade tissues are consistent with characteristics associated with acclimation to higher light, resembling those of sun leaves. Previous studies by Terashima and Inoue (1984, 1985a, b) have shown that photosynthetic properties within a leaf exhibit a vertical gradient from sun-type characteristics on the adaxial side to shade-type characteristics toward the abaxial side. The PAM measurements of the present study are consistent with this concept and further show that this sun–shade-like differentiation within a leaf is also reflected in dynamic photochemical responses.

### Relatively uniform tissue in rice leaves

In rice, the leaf mesophyll tissues do not differentiate into distinct palisade and spongy tissues. Instead, the mesophyll tissue is composed of lobed mesophyll cells distributed throughout the leaf cross-section. Therefore, we divided the mesophyll into upper and lower regions along the adaxial–abaxial axis for analysis. Both Y(II) and Y(NPQ) increased similarly in the upper and lower regions of the leaf, and Y(NO) also decreased similarly (Fig. 5, Supplementary Movie 4). There was no significant difference in F_v_/F_m_ between the upper and lower tissues (Table 1).

**Fig. 5.**
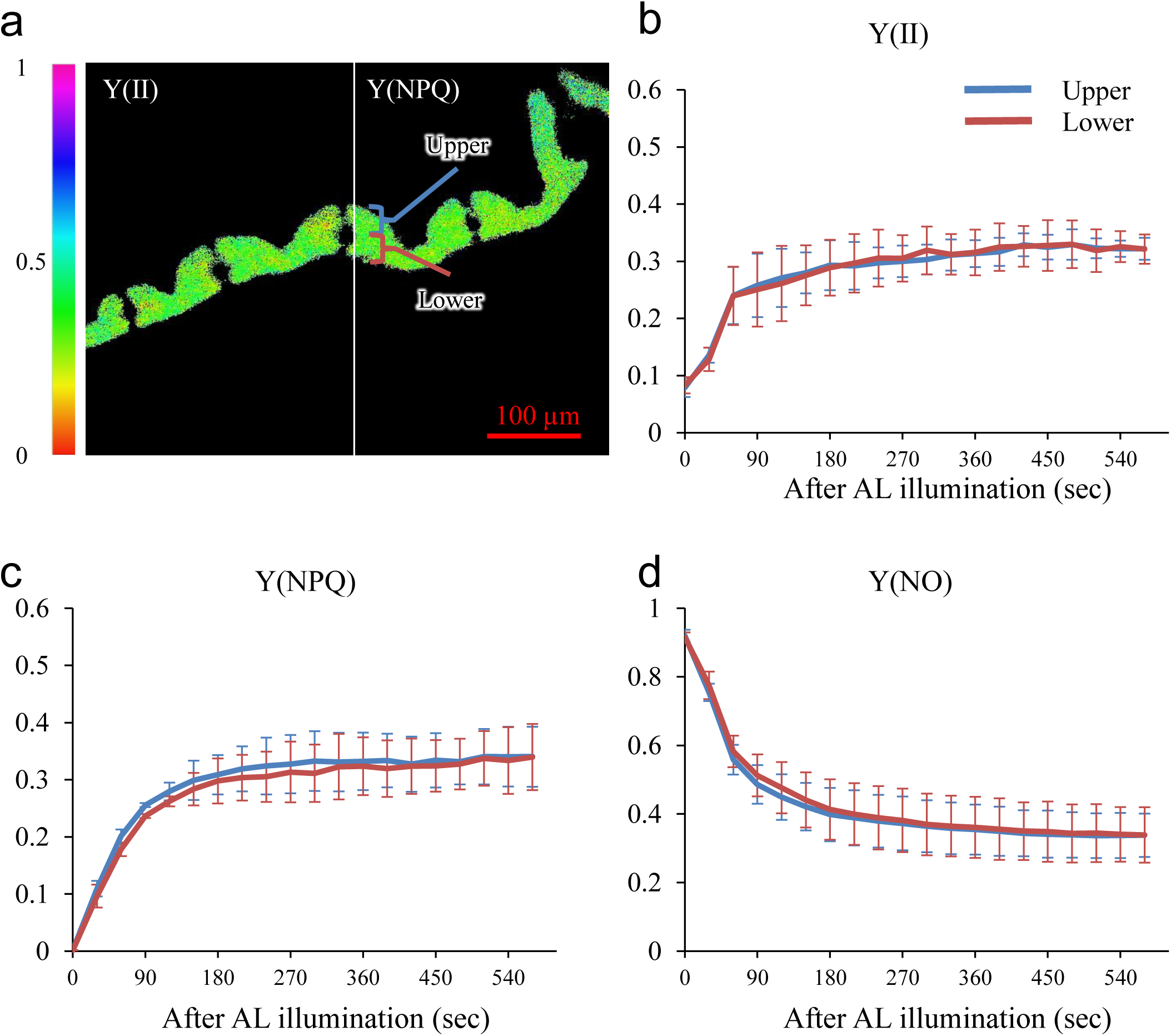
Microscopic Imaging-PAM analysis of rice leaf transverse sections. (a) Y(II) (left side) and Y(NPQ) (right side) images of a transverse leaf section of rice acquired using microscopic Imaging-PAM at the end of the induction period. The upper side corresponds to the adaxial side of the leaf. (b–d) Induction curves of Y(II), Y(NPQ), and Y(NO) measured in upper and lower regions. Values are means ± SD. n = 3.

Rice mesophyll tissues consist of highly lobed cells that form a complex three-dimensional geometry across the leaf thickness (Chonan 1967; Oi et al. 2017; Ouk et al. 2022). A 3D reaction-diffusion model incorporating ray tracing of light propagation based on the reconstructed anatomy of rice leaves has estimated that an increase in the number of lobes would lead to more uniform light absorption within the leaf (Xiao et al., 2023). Furthermore, although incident light generally enters leaves from the adaxial side, the erect leaf posture typical of grasses such as rice is thought to allow light to enter from the abaxial side more readily than in the more horizontally oriented leaves of dicots. As a result, the light environment across the adaxial–abaxial axis becomes more balanced. Together, these structural features are likely to reduce the formation of steep internal light gradients. This may explain why rice leaves exhibited little spatial differentiation in Y(II) and Y(NPQ), suggesting that photosynthetic regulation operates more uniformly across the leaf. However, it should be noted that unilateral illumination does not produce a completely uniform internal light distribution even in rice leaves (Takahashi et al., 1994). Indeed, PAM measurements performed separately from the adaxial and abaxial sides of rice flag leaves have revealed measurable differences in photosynthetic parameters (Kumagai et al. 2014). Therefore, the anatomy of rice leaves does not completely equalize the light environment under all conditions but rather facilitates a tendency toward greater uniformity compared with leaves that possess clear palisade–spongy differentiation.

### Mesophyll and bundle sheath cells in C_4_ leaves

In finger millet, mesophyll and bundle sheath cells were analyzed separately. Y(II) increased rapidly in both mesophyll (approximately 0.3) and bundle sheath cells (approximately 0.4) within the first 30 s. Thereafter, it gradually declined, stabilizing at approximately 0.2 in mesophyll cells and approximately 0.3 in bundle sheath cells (Fig. 6, Supplementary Movie 5). Y(NPQ) increased gradually in both mesophyll and bundle sheath cells and reached approximately 0.35 in mesophyll cells and approximately 0.25 in bundle sheath cells at approximately 240 s after AL illumination. Y(NO) exhibited a similarly rapid decrease in mesophyll and bundle sheath cells. There was no significant difference in F_v_/F_m_ between mesophyll and bundle sheath cells (Table 1).

**Fig. 6.**
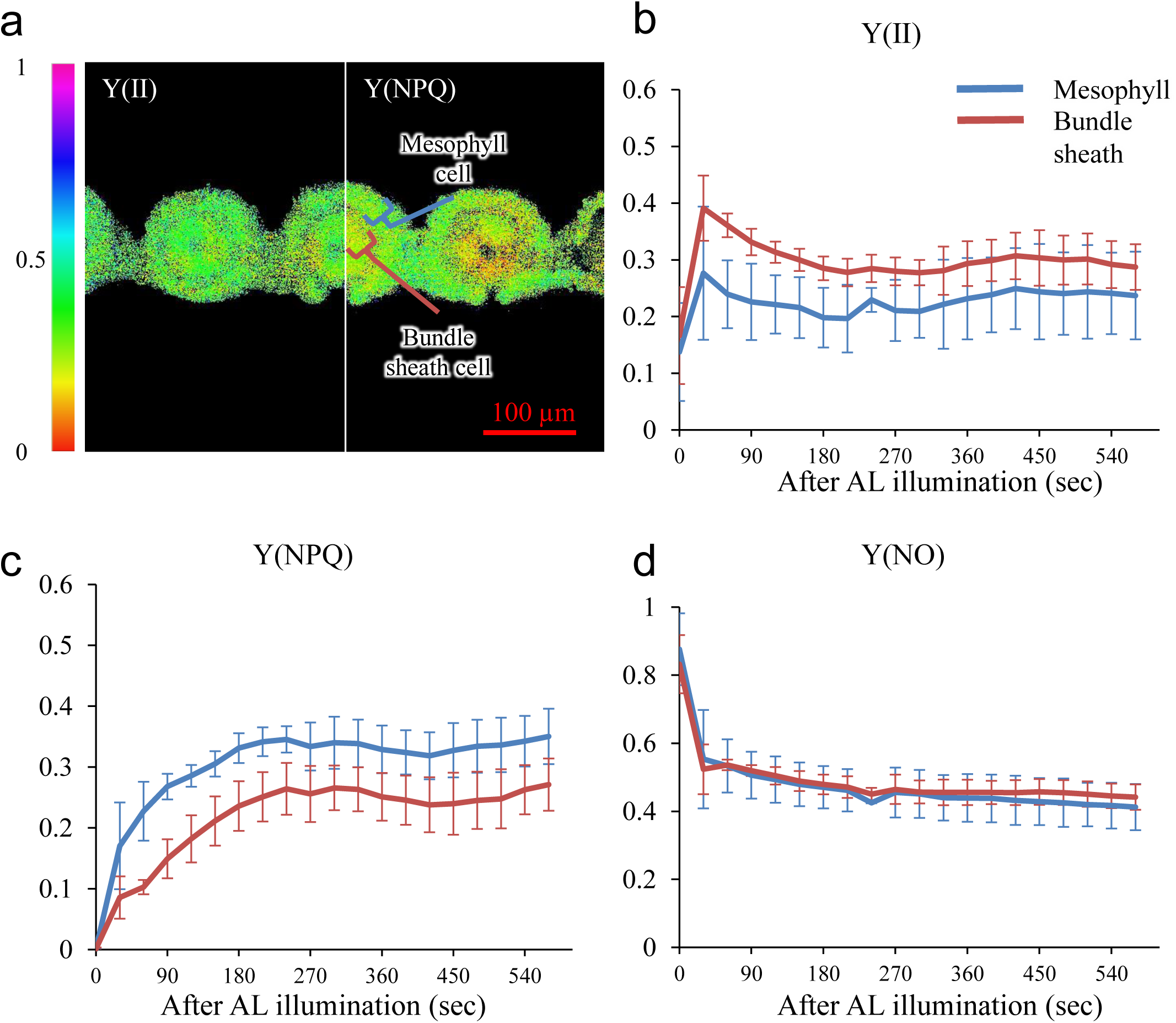
Microscopic Imaging-PAM analysis of finger millet leaf transverse sections. (a) Y(II) (left side) and Y(NPQ) (right side) images of a transverse leaf section of finger millet acquired using microscopic Imaging-PAM at the end of the induction period. The upper side corresponds to the adaxial side of the leaf. (b–d) Induction curves of Y(II), Y(NPQ), and Y(NO) measured in mesophyll and bundle sheath regions. Values are means ± SD. n = 4.

C_4_ plants exhibit functional differentiation between mesophyll and bundle sheath cells, with phosphoenolpyruvate carboxylase localized mainly in mesophyll cells and Rubisco localized in bundle sheath cells (Dengler and Nelson 1999). The abundance and organization of the photosystems also differ between the two cell types. Bundle sheath chloroplasts generally contain fewer PSII complexes than mesophyll chloroplasts (Ghannoum et al. 2005). Despite this lower PSII abundance, Y(II) remained relatively high in bundle sheath cells in finger millet (Fig. 6B). This pattern may reflect differences in the balance between the generation and consumption of reducing power through photosynthetic electron transport. In C_4_ photosynthesis, the Calvin cycle operates in bundle sheath cells and requires reducing equivalents. Accordingly, linear electron transport in bundle sheath chloroplasts is expected to play an important role in supplying reducing power for the Calvin cycle, although intercellular exchange of metabolites (e.g., via the 3-PGA/triose-phosphate shuttle) may also contribute. As a result, reducing power generated in bundle sheath chloroplasts may be efficiently consumed by the Calvin cycle, which may help sustain relatively high Y(II) in these cells. In contrast, mesophyll cells possess a comparatively weaker sink for reducing equivalents because the Calvin cycle is absent in these cells. Consequently, excess excitation energy may be more readily dissipated through non-photochemical quenching, resulting in higher Y(NPQ).

Interestingly, an opposite tendency has been reported in NADP-ME-type C_4_ species, including maize, compared with the pattern observed here in the NAD-ME-type species finger millet (Ghannoum et al. 2005; Liu et al. 2022). In maize, F_v_/F_m_ and Y(II) tend to be higher in mesophyll cells, whereas NPQ is higher in bundle sheath cells. In NADP-ME plants, malate is the major C_4_ acid transported between mesophyll and bundle sheath cells, and malate decarboxylation generates NADPH in bundle sheath chloroplasts (Kanai and Edwards 1999). This additional source of reducing power may decrease the requirement for linear electron transport in bundle sheath cells, which may lead to lower Y(II) and increased dissipation of excitation energy as NPQ. These results suggest that the balance between photochemical electron transport and non-photochemical energy dissipation differs between mesophyll and bundle sheath cells in C_4_ leaves, and that these cell-type-specific fluorescence responses may vary among C_4_ biochemical subtypes.

Recognizing the differences between the section-based approach of the present study and conventional whole-leaf measurement methods is critically important. The most fundamental distinction lies in how light propagates within the leaf. In intact leaves, incident light enters through the epidermis and is redistributed within the tissue through repeated refraction and scattering at air–liquid interfaces in intercellular spaces, as well as through absorption and transmission by individual chloroplasts (Terashima and Saeki, 1983; Terashima et al., 2009). As a result, internal light gradients are strongly shaped by leaf anatomy and optical properties. In contrast, in our sectioning approach, intercellular air spaces were removed by infiltration, and illumination was applied directly to the surface of the sections. This configuration largely eliminates the complex internal light gradients present in intact leaves and enables more uniform illumination across different tissues. In addition, the absolute photon flux experienced by leaf tissues in cross-sections is likely to differ substantially from that in intact leaves due to the fundamentally different light paths. Therefore, even when nominal light settings are identical, direct quantitative comparisons between intact-leaf and section-based measurements are not straightforward. Moreover, sectioning and infiltration simultaneously alter CO₂ diffusion, water status, ionic balance, and oxygen conditions, potentially disturbing native leaf physiology. The F_v_/F_m_ values obtained in the present study (approximately 0.5–0.7; Table 1) were lower than the values of around 0.8 typically observed in healthy, dark-adapted leaves. These relatively low values may reflect both physiological effects of section preparation and technical limitations associated with microscopic fluorescence imaging. In particular, accurate determination of F_o_ becomes challenging at microscopic scales because of the reduced fluorescence signal from small detection areas; overestimation of F_o_ would directly lower the calculated F_v_/F_m_ (Oxborough 2004). Importantly, although our approach does not reproduce the native optical and physiological environment of intact leaves, it creates a simplified and more uniform light condition across tissues. Under these conditions, differences in photochemical behavior cannot be attributed to internal light gradients, but instead reflect intrinsic, tissue-specific properties. Consequently, the section-based approach provides a powerful complementary framework for revealing tissue-dependent heterogeneity in photosynthetic regulation that is otherwise masked by the complex light environment of intact leaves.

The results of the present study reveal that leaves are not uniform photosynthetic organs but are composed of tissues that exhibit distinct photochemical and photoprotective behaviors depending on their anatomical context. Although the present study examined photosynthetic characteristics in fully expanded leaves of several species under short-term illumination, this approach also provides opportunities to investigate temporal changes and responses to various environmental stresses at the tissue level. By applying this method to leaves subjected to different acclimation histories or stress treatments, it becomes possible to assess how photochemical responses are spatially redistributed within leaves. Indeed, using the same method, we have observed that low-temperature treatment in cold-sensitive cucumber preferentially reduces Y(II) in palisade tissues, indicating that stress responses may occur in a tissue-specific manner within leaves (Takeuchi and Kato et al. in preparation; Takeuchi et al., 2025). By enabling tissue-resolved chlorophyll fluorescence measurements in living leaves across different plant types, this approach provides a new framework for understanding how leaf architecture and photosynthetic strategy shape the spatial organization of light utilization within leaves.

## Materials and Methods

### Plant material and growth conditions

Cucumber (*Cucumis sativus* cv. ‘High Green 21’) seeds were purchased from Saitama Gen-shu Ikuseikai (Saitama, Japan). Finger millet (*Eleusine coracana*) seeds were purchased from Snow Brand Seed Co., Ltd. (Hokkaido, Japan). Maize (*Zea mays* cv. Honey Bantam 20) seeds were purchased from Sakata Seed Co., Ltd. (Kanagawa, Japan). Cucumber, finger millet, maize, and Arabidopsis (*Arabidopsis thaliana*, ecotype Columbia-0) were grown in plastic pots filled with soil in a growth chamber under a 16 h light / 8 h dark photoperiod at 27 °C with a light intensity of 250–300 μmol photons m⁻² s⁻¹. Fully expanded upper leaves from 2–3-week-old plants were used for analysis.

Rice (*Oryza sativa* japonica cv. ‘Hitomebore’) seeds were soaked in distilled water containing fungicide (Trifumin; Nippon Soda, Tokyo, Japan) for 2 days, then transferred to a hydroponic medium and grown under the same environmental conditions described above. The hydroponic medium consisted of the following components in tap water (Takeuchi et al., 2024): 0.5 mM CaCl_2_, 0.25 mM KH_2_PO4, 1 mM (NH_4_)_2_SO_4_, 0.5 mM MgCl_2_, 0.5 mM KCl, 2.86 mg L^-1^ H_3_BO_3_, 1.81 mg L^-1^ MnCl_2_·4H_2_O, 0.08 mg L^-1^ CuSO_4_·5H_2_O, 0.22 mg L^-1^ ZnSO_4_·7H_2_O, 0.1 mg L^-1^ (NH_4_)_6_Mo_7_O_24_·4H_2_O, and 90 μM Fe (III)-EDTA. The medium was refreshed twice a week. The highest fully expanded leaves from approximately 1-month-old plants were used for analysis.

Camellia (*Camellia japonica*) young plants approximately 50 cm in height were purchased from a commercial nursery and acclimated for approximately one month in a glasshouse under natural light conditions (temperature: 22–26 °C) prior to use in the experiments.

### Preparation of live leaf cross-sections

Live leaf sections were prepared as previously described (Kato et al., 2025b), with minor modifications. Leaves were excised and trimmed into approximately 5 × 5 mm segments. The leaf segments were infiltrated with mounting solution containing 50 mM MES-NaOH (pH 6.0), 2 mM NaHCO_3_, 1 mM CaCl_2_, and 1 mM MgCl_2_ using a syringe to remove air from the intercellular spaces. The infiltrated leaf segments were manually sectioned using a razor blade. A silicone rubber sheet was cut into a square frame and pasted to a glass slide to create a groove (Fig. 1A, B). Mounting solution was added to the groove, and a leaf section was transferred into the solution. The sample was then covered with a cover glass to seal the section in the mounting medium. Rice leaves were infiltrated after sectioning because the small intercellular spaces made infiltration of the leaf segments difficult. Detailed procedures for live leaf-section preparation are described in Kato et al. (2025b).

### Microscopic Imaging-PAM

Chlorophyll fluorescence in leaf tissues was measured at room temperature (approximately 25 °C) using a Walz IMAGING-PAM M-Series chlorophyll fluorescence imaging system (Heinz Walz GmbH, Effeltrich, Germany), equipped with a blue LED excitation module (IMAG-L470M). Leaf sections mounted on the microscopic Imaging-PAM stage were first observed under bright-field illumination to confirm that section thickness was comparable among samples and that tissue structure was not collapsed as a result of mechanical contact during the procedures. After the position of the section was adjusted, the microscope light was turned off, and the samples were dark-adapted for 15 min. Chlorophyll fluorescence induction during the dark-to-light transition was then monitored for 9.5 min using the induction-curve mode of the Imaging-PAM system. The measuring light (ML) and actinic light (AL) intensities were set to ML 2 (PPFD 2.2 µmol m⁻² s⁻¹) and AL 4 (PPFD 115.8 µmol m⁻² s⁻¹), respectively. Saturating pulses (PPFD ≈ 4,000 µmol m⁻² s⁻¹) were applied at 30-s intervals throughout the measurement. The light intensities at the sample plane were measured using an MC-MQS/OVV microscopy quantum sensor (Heinz Walz GmbH). The maximum quantum yield of PSII after dark adaptation was calculated as F_v_/F_m_ = (F_m_ − Fo)/F_m_. The effective quantum yield of PSII was calculated as Y(II) (ΦII) = (F’_m_ − F_s_)/F*’_m_*. The quantum yields of nonregulated and regulated energy dissipation in PSII were calculated as Y(NO) (ΦNO) = F_s_/F_m_ and Y(NPQ) (ΦNPQ) = F_s_/F’_m_ − F_s_/F_m_, respectively. ‘Y’ corresponds to quantum yields and is equivalent to Φ used in classical photochemistry. The F_o_ and F_m_ levels, the minimum and maximum fluorescence, respectively, were measured after leaves were kept in the dark for 15 min. During the induction-curve measurement, F’_m_ (maximum fluorescence under AL) and F_s_ (steady-state fluorescence under light) were measured.

Areas of interest (AOIs) were defined for each tissue type, and three AOIs were selected per tissue within each section (Supplementary Fig. S1). The mean of these three AOIs in a section was treated as a single biological replicate. In *Arabidopsis* and cucumber, AOIs were separately assigned to palisade and spongy mesophyll tissues (Supplementary Fig. S1A, B). In Camellia leaves, the palisade mesophyll was further subdivided into three layers according to cell layers, and the spongy mesophyll was similarly divided into three vertical regions (Supplementary Fig. S1C). In rice leaves, the mesophyll was divided into upper and lower regions along the adaxial–abaxial axis (Supplementary Fig. S1D). In the C_4_ species, AOIs were defined separately for mesophyll and bundle sheath cells (Supplementary Fig. S1E, F).

### Statistical analysis

Data were analyzed using Microsoft Excel with the add-in software package Statcel 3 (OMS Publishing, Japan). Statistical analyses were based on Student’s *t*-test or the Tukey–Kramer method. Details of statistical parameters are provided in the figure legends.

## Acknowledgements

The authors thank Associate Professor Wataru Yamori (The University of Tokyo) for kindly providing access to the imaging-PAM system and for valuable advice on this study.

During the preparation of this manuscript, the authors used ChatGPT (OpenAI) to assist with language editing and wording. All suggestions were reviewed and revised by the authors, who take full responsibility for the final content.

## Author contributions

**Y.K.** conceived the study, performed all experiments, and wrote the manuscript. **K.T.** and **S.H.** assisted with the experiments and contributed to data interpretation. **K.K.** assisted with the use of the Imaging-PAM system and provided technical guidance for the experiments. **K.I.** supervised the project and contributed to the development of the research plan. All authors reviewed and approved the final manuscript.

## Data Availability

The data underlying this article are available from the corresponding author upon reasonable request.

