## Supplemental Figure for "Tissue-resolved photosynthetic responses in living leaves revealed by microscopic imaging-pulse-amplitude-modulation"

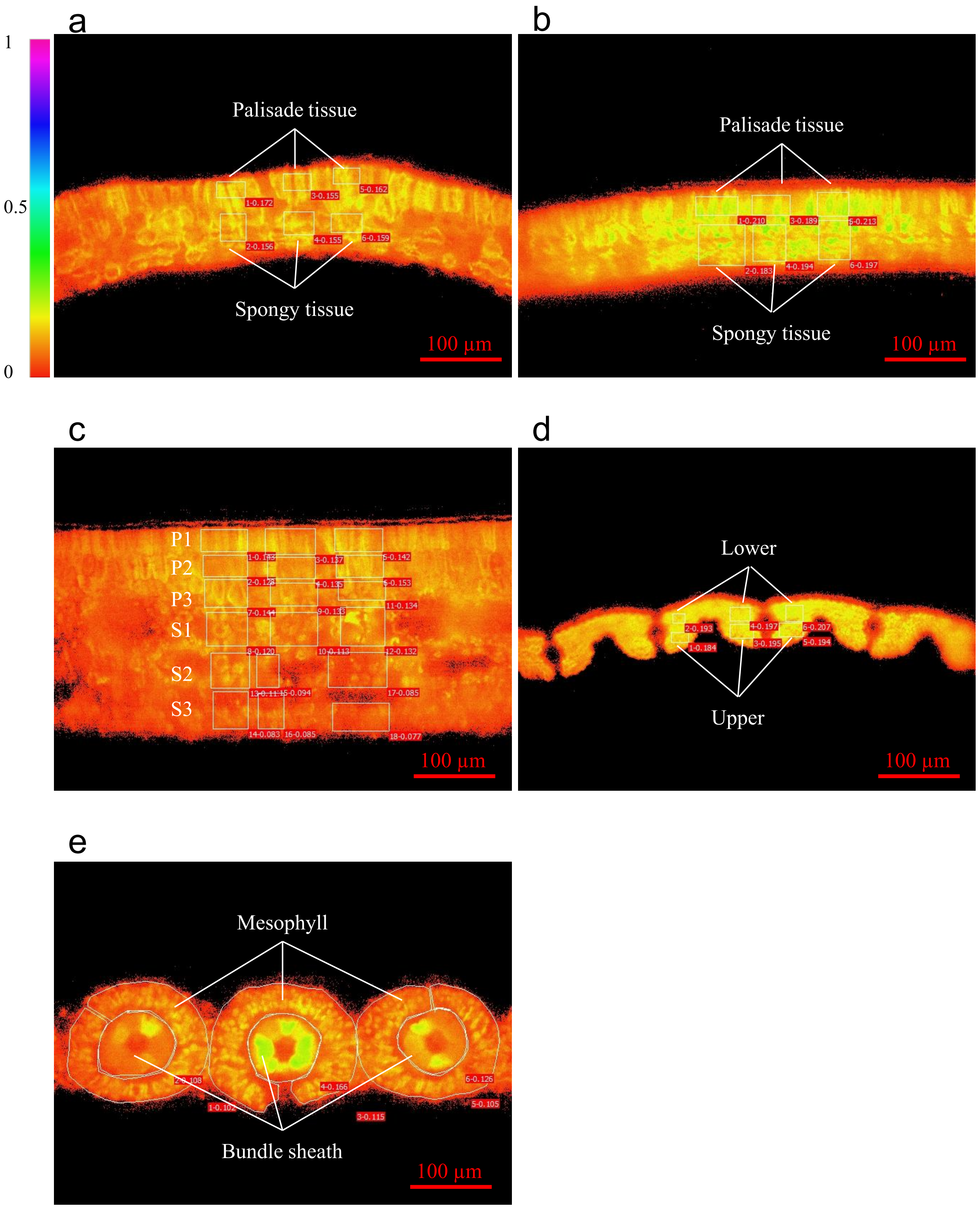


**Supplementary Fig. S1 AOI (area of interest) settings for each plant species.**
Fm microscopic images of leaf transverse sections acquired using microscopic Imaging-PAM. Each white outline indicates a single AOI, and the numbers shown within the red boxes represent the mean Fm value for that AOI. The upper side of the image corresponds to the adaxial side in all species except rice. In rice, the lower side of the image corresponds to the adaxial side.
(a) Arabidopsis, (b) cucumber, (c) camellia, (d) rice, and (e) finger millet.

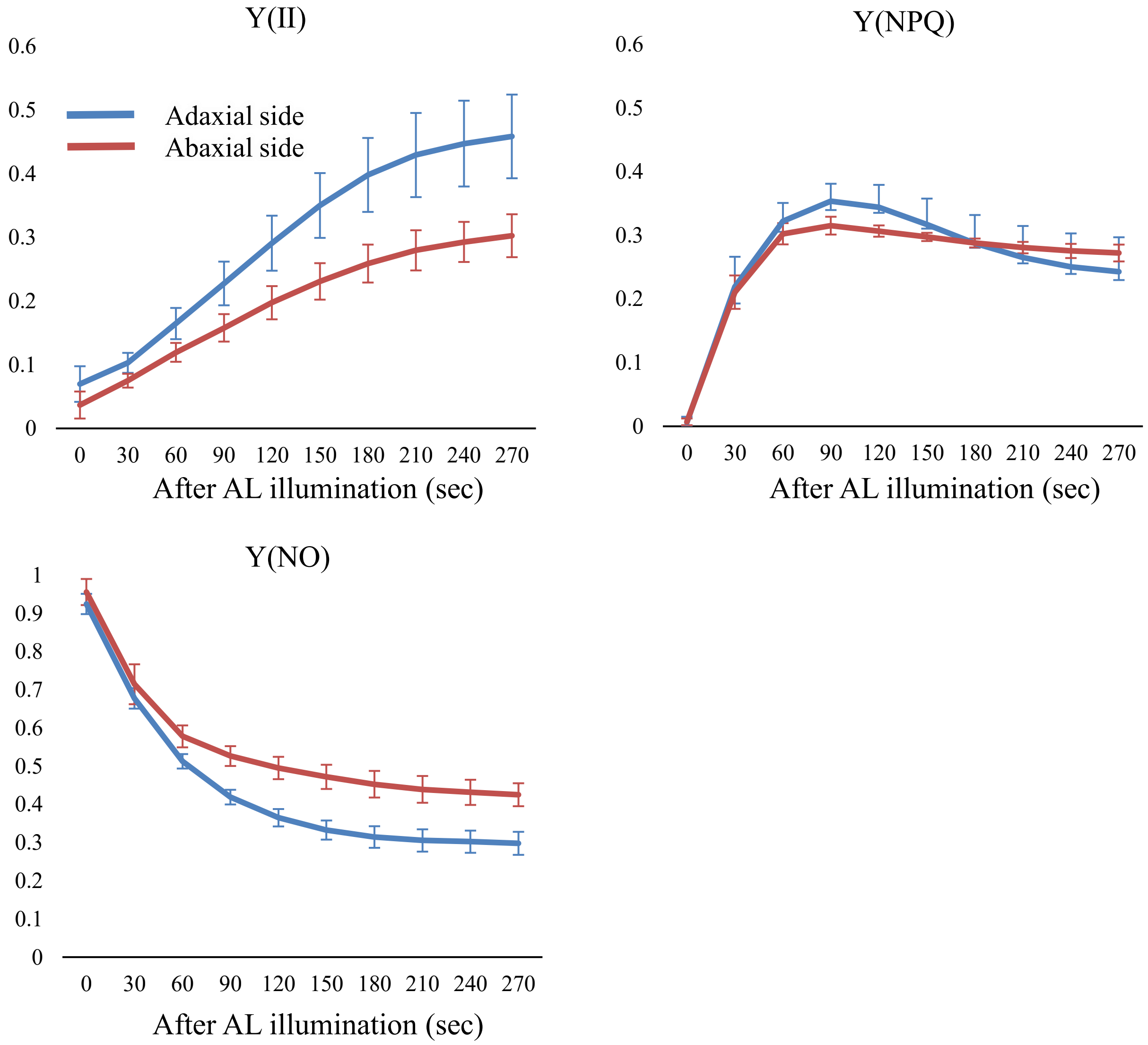


**Supplementary Fig. S2 PAM analysis of cucumber leaf from the adaxial or abaxial side.**
(a-c) Induction curves of Y(II), Y(NPQ), and Y(NO) measured from the adaxial or abaxial side. Values are means ± SD. n = 3. Chlorophyll fluorescence parameters were measured using a Dual-PAM 100 (Heinz Walz GmbH, Effeltrich, Germany).

Supplementary Table 1. $F_v/F_m$ value measured from adaxial or abaxial side.

| | **Adaxial** | **Abaxial** | |
|---|---|---|---|
| | 0.82±0.01 | 0.80±0.01 | * (*p*= 0.01) |

Each value represents the mean (±SD). Asterisk indicate a significant difference at $p < 0.05$ (Paired t-test, n = 3.)
